# European Primary Forest Database (EPFD) v2.0

**DOI:** 10.1101/2020.10.30.362434

**Authors:** Francesco Maria Sabatini, Hendrik Bluhm, Zoltan Kun, Dmitry Aksenov, José A. Atauri, Erik Buchwald, Sabina Burrascano, Eugénie Cateau, Abdulla Diku, Inês Marques Duarte, Ángel B. Fernández López, Matteo Garbarino, Nikolaos Grigoriadis, Ferenc Horváth, Srđan Keren, Mara Kitenberga, Alen Kiš, Ann Kraut, Pierre L. Ibisch, Laurent Larrieu, Fabio Lombardi, Bratislav Matovic, Radu Nicolae Melu, Peter Meyer, Rein Midteng, Stjepan Mikac, Martin Mikoláš, Gintautas Mozgeris, Momchil Panayotov, Rok Pisek, Leónia Nunes, Alejandro Ruete, Matthias Schickhofer, Bojan Simovski, Jonas Stillhard, Dejan Stojanovic, Jerzy Szwagrzyk, Olli-Pekka Tikkanen, Elvin Toromani, Roman Volosyanchuk, Tomáš Vrška, Marcus Waldherr, Maxim Yermokhin, Tzvetan Zlatanov, Asiya Zagidullina, Tobias Kuemmerle

## Abstract

Primary forests are scarce in Europe and continue to disappear at an alarming rate. Despite these losses, we know little about where such forests still occur. Here, we present an updated geodatabase and map of Europe’s known primary forests. Our geodatabase harmonizes 48 different datasets of primary forests, and contains 18,411 individual patches (41.1 Mha) spread across 33 countries. When available, we provide information on each patch (name, location, naturalness, extent and dominant tree species) and the surrounding landscape (biogeographical regions, protection status, potential natural vegetation, current forest extent). To assess the robustness of our geodatabase, we checked each patch for forest disturbance events using Landsat satellite-image time series (1985-2018). We estimate that 94% of the patches in our database did not experience significant disturbances that would alter their primary forest status in the last 30 year. Our database is the most comprehensive dataset on primary forests in Europe, and will be useful for biogeographic and ecological studies, and conservation planning to safeguard these unique forests.

## Background & Summary

Primary forests are composed of native tree species without clearly visible indications of human activity and with intact ecological processes^1,2^. The importance of such forests is widely recognized^3,4^. First, they provide refuge to forest biodiversity^5^, and act as a buffer to species loss in human-dominated landscapes^6^. Second, primary forests play an important role in climate change mitigation. At the local scale, they buffer the adverse effects of increasing temperature on understory biodiversity, as they often have cooler forest-floor summer temperatures compared to secondary forests^7^. At the global scale they contribute to climate stability by storing large quantities of carbon, both in the biomass and in soils^3,8,9^. Third, primary forests often serve as a reference for developing close-to-nature forest management, or for benchmarking restoration efforts^10^. Finally, these forests are an irreplaceable part of our natural heritage, shape the cultural identities of local communities, and have a high intrinsic value^11^.

In Europe, as in many human-dominated regions, most forested area is currently managed^12^, often with increasing harvest intensities^13,14^. As a result, despite the general trend of increasing total forest area, primary forests are scarce and continue to disappear^15^. For instance, Romania hosts some of the largest swaths of primary forest in Central Europe and faced a sharp increase in logging rates since 2000. This has resulted in significant primary forest loss, even within protected areas^15–17^. In Poland, the iconic Białowieża Forest was recently in the spotlight after the controversial decision from the Polish National Forest Holding, now nullified by the Court of Justice of the European Union^18^, to implement salvage logging followed by tree planting after a bark beetle outbreak^19^. Widespread loss of primary forests also occurred in Ukraine^20^, Slovakia^21^, or in the boreal North, e.g., in the Russian North-West, where 4.6 Mha of primary forest were lost since 2001^15,22^. Effective protection of Europe’s primary forests is therefore urgently needed^23^.

In the newly released ‘Biodiversity Strategy for 2030’, the European Commission emphasized the need to define, map, monitor and strictly protect all of the EU’s remaining primary and old-growth forests^4^. Reaching these objectives requires complete and up-to-date data on primary forests’ location and protection status. Such data could inform both conservation planning and research, for instance by highlighting areas where primary forests are either scarce, or poorly studied. Yet, many data gaps remain on the location and conservation status of EU’s primary forests^23^. Only a few countries conducted systematic, on-the-ground inventories^21,24^. For most countries data are either only available for a few well-studied forests^25–27^, or are limited to the distribution of potential (=unconfirmed) primary forests, typically predicted statistically or via remote sensing^28–30^. Despite past efforts for harmonizing data^31,32^, only recently has the first map of primary forests been released for Europe^33^ together with a first assessment of their conservation status^23^.

The first version of our European Primary Forest database (EPFD v1.0) included 32 local-to-national datasets, plus data from a literature review and a survey, resulting in the mapping of a total of ~1.4 Mha of primary forest^33^. This is only about one fifth of the estimated 7.3 Mha of undisturbed forest still occurring in Europe, excluding Russia^12^. Here, we build on those efforts to substantially progress towards a complete EPFD, as well as to release the data open-access^34^. Key improvements of this new database include (a) filling major regional gaps, including European Russia, the Balkan Peninsula, the Pyrenees and the Baltic region, (2) mapping ‘potential’ primary forests for Sweden and Norway, two key regions where complete inventories are currently unavailable, and (3) updating our literature review to January 2019.

EPFD v2.0 thus aggregates and harmonizes 48 regional-to-continental spatial datasets, contains 18,411 non-overlapping primary forest patches (plus 299 point features) covering an area of 41.1 Mha (37.4 Mha in European Russia alone; Figure 1) across 33 countries (Table 1). Potential primary forests for Sweden and Norway account for an additional 16,311 polygons and 2.5 Mha (Figure 2).

**Figure 1.**
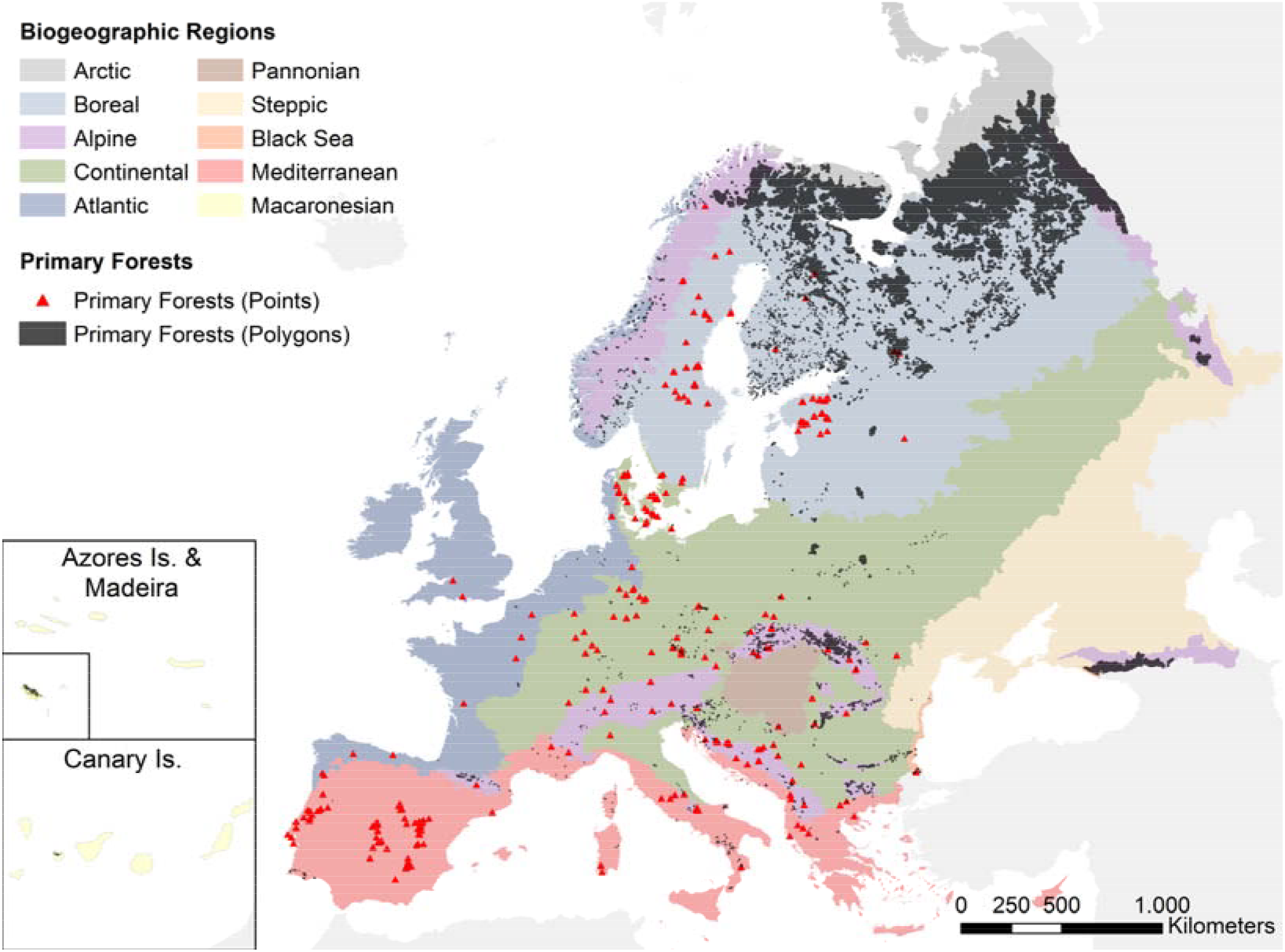
Overview of the primary forest patches contained in the EPFD v2.0. Both points and polygons were magnified to improve visibility.

**Table 1.**
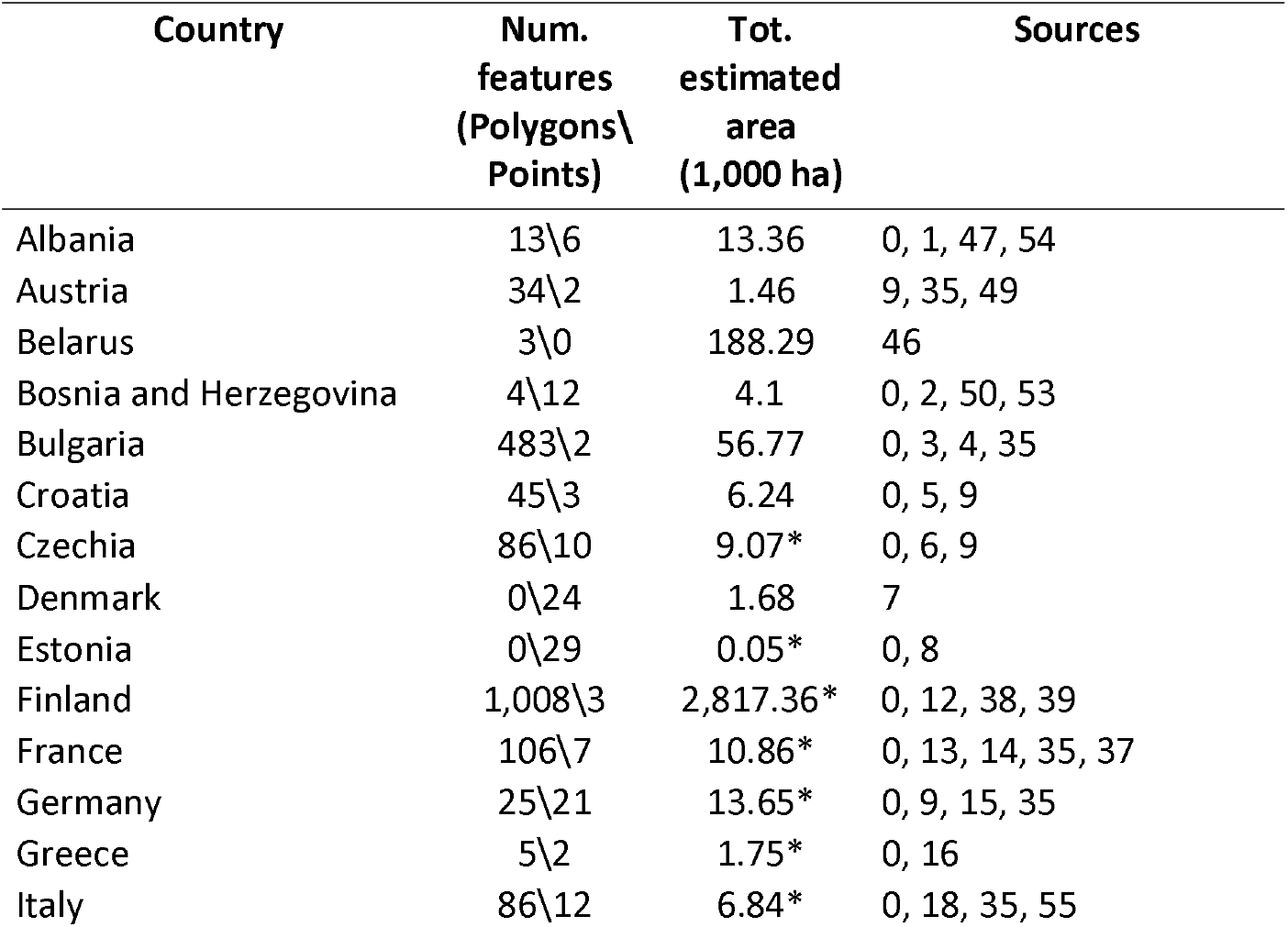
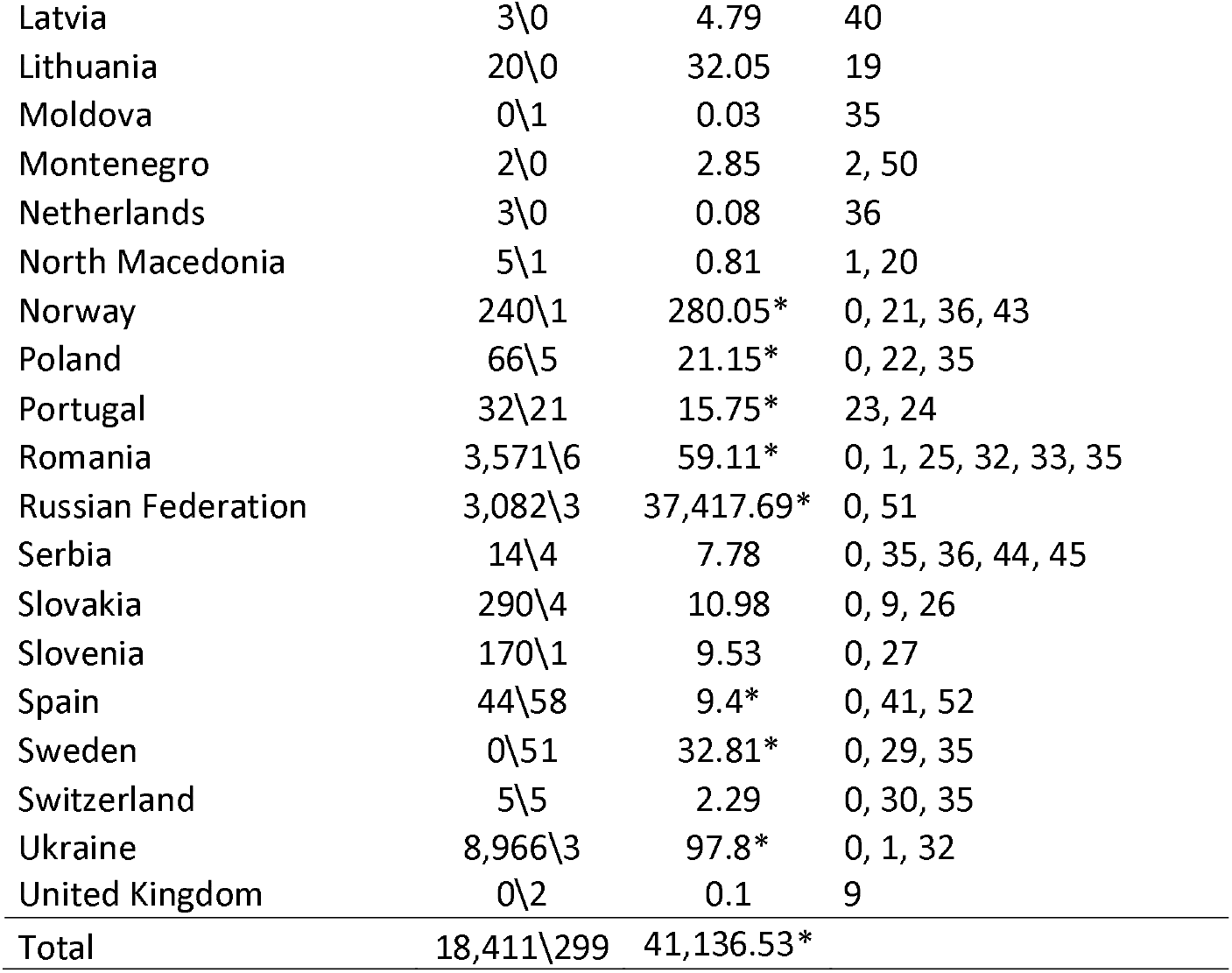
Summary of primary forest data across European countries. Dataset IDs correspond to those in Table 2. * Some point features have no information on forest patch area.

| Country | Num.<br>features<br>(Polygons\<br>Points) | Tot.<br>estimated<br>area<br>(1,000 ha) | Sources |
| --- | --- | --- | --- |
| Albania | 13\6 | 13.36 | 0, 1, 47, 54 |
| Austria | 34\2 | 1.46 | 9, 35, 49 |
| Belarus | 3\0 | 188.29 | 46 |
| Bosnia and Herzegovina | 4\12 | 4.1 | 0, 2, 50, 53 |
| Bulgaria | 483\2 | 56.77 | 0, 3, 4, 35 |
| Croatia | 45\3 | 6.24 | 0, 5, 9 |
| Czechia | 86\10 | 9.07* | 0, 6, 9 |
| Denmark | 0\24 | 1.68 | 7 |
| Estonia | 0\29 | 0.05* | 0, 8 |
| Finland | 1,008\3 | 2,817.36* | 0, 12, 38, 39 |
| France | 106\7 | 10.86* | 0, 13, 14, 35, 37 |
| Germany | 25\21 | 13.65* | 0, 9, 15, 35 |
| Greece | 5\2 | 1.75* | 0, 16 |
| Italy | 86\12 | 6.84* | 0, 18, 35, 55 |
| Latvia | 3\0 | 4.79 | 40 |
| Lithuania | 20\0 | 32.05 | 19 |
| Moldova | 0\1 | 0.03 | 35 |
| Montenegro | 2\0 | 2.85 | 2, 50 |
| Netherlands | 3\0 | 0.08 | 36 |
| North Macedonia | 5\1 | 0.81 | 1, 20 |
| Norway | 240\1 | 280.05* | 0, 21, 36, 43 |
| Poland | 66\5 | 21.15* | 0, 22, 35 |
| Portugal | 32\21 | 15.75* | 23, 24 |
| Romania | 3,571\6 | 59.11* | 0, 1, 25, 32, 33, 35 |
| Russian Federation | 3,082\3 | 37,417.69* | 0, 51 |
| Serbia | 14\4 | 7.78 | 0, 35, 36, 44, 45 |
| Slovakia | 290\4 | 10.98 | 0, 9, 26 |
| Slovenia | 170\1 | 9.53 | 0, 27 |
| Spain | 44\58 | 9.4* | 0, 41, 52 |
| Sweden | 0\51 | 32.81* | 0, 29, 35 |
| Switzerland | 5\5 | 2.29 | 0, 30, 35 |
| Ukraine | 8,966\3 | 97.8* | 0, 1, 32 |
| United Kingdom | 0\2 | 0.1 | 9 |
| Total | 18,411\299 | 41,136.53* |  |

**Table 2.**
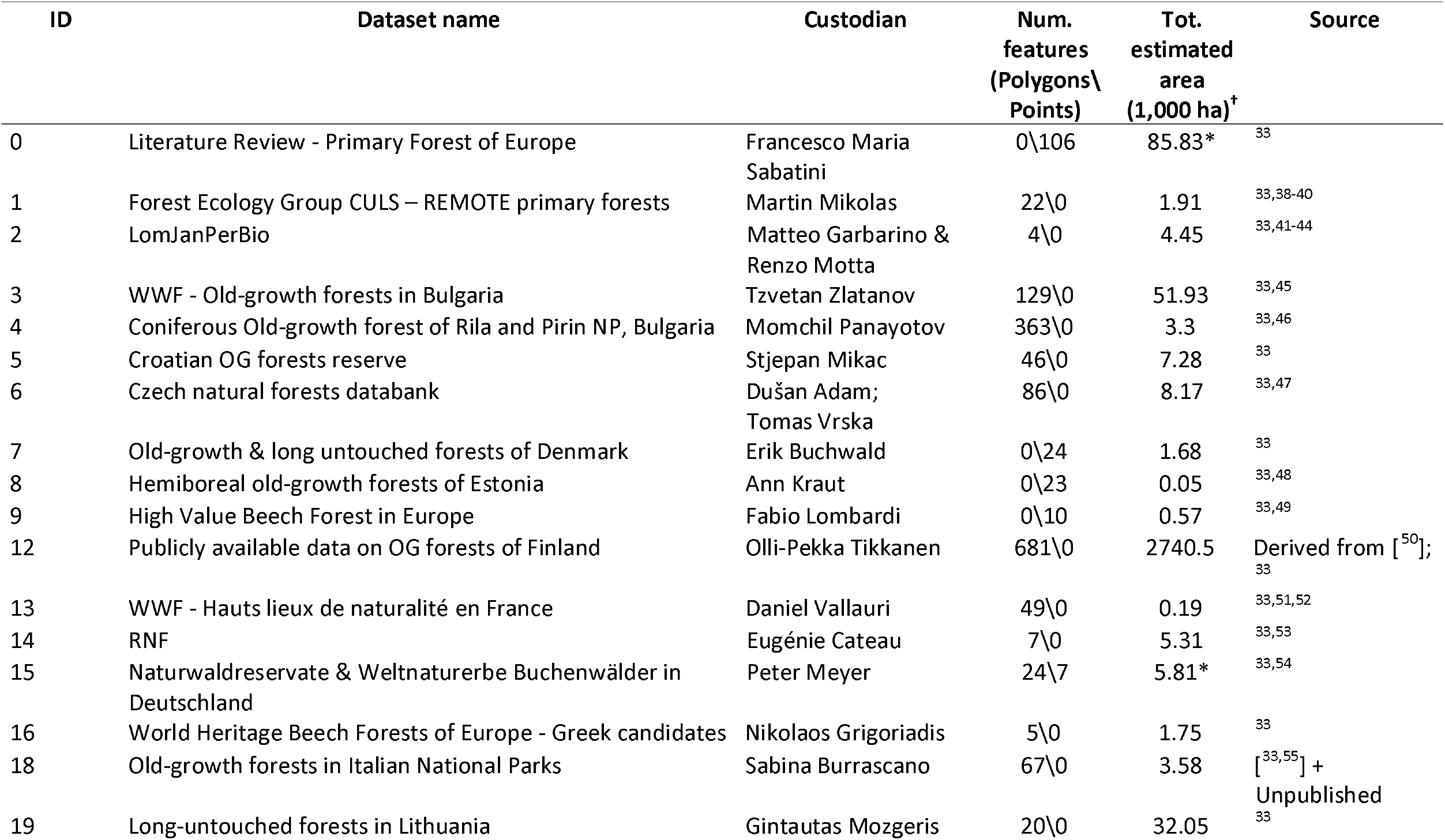
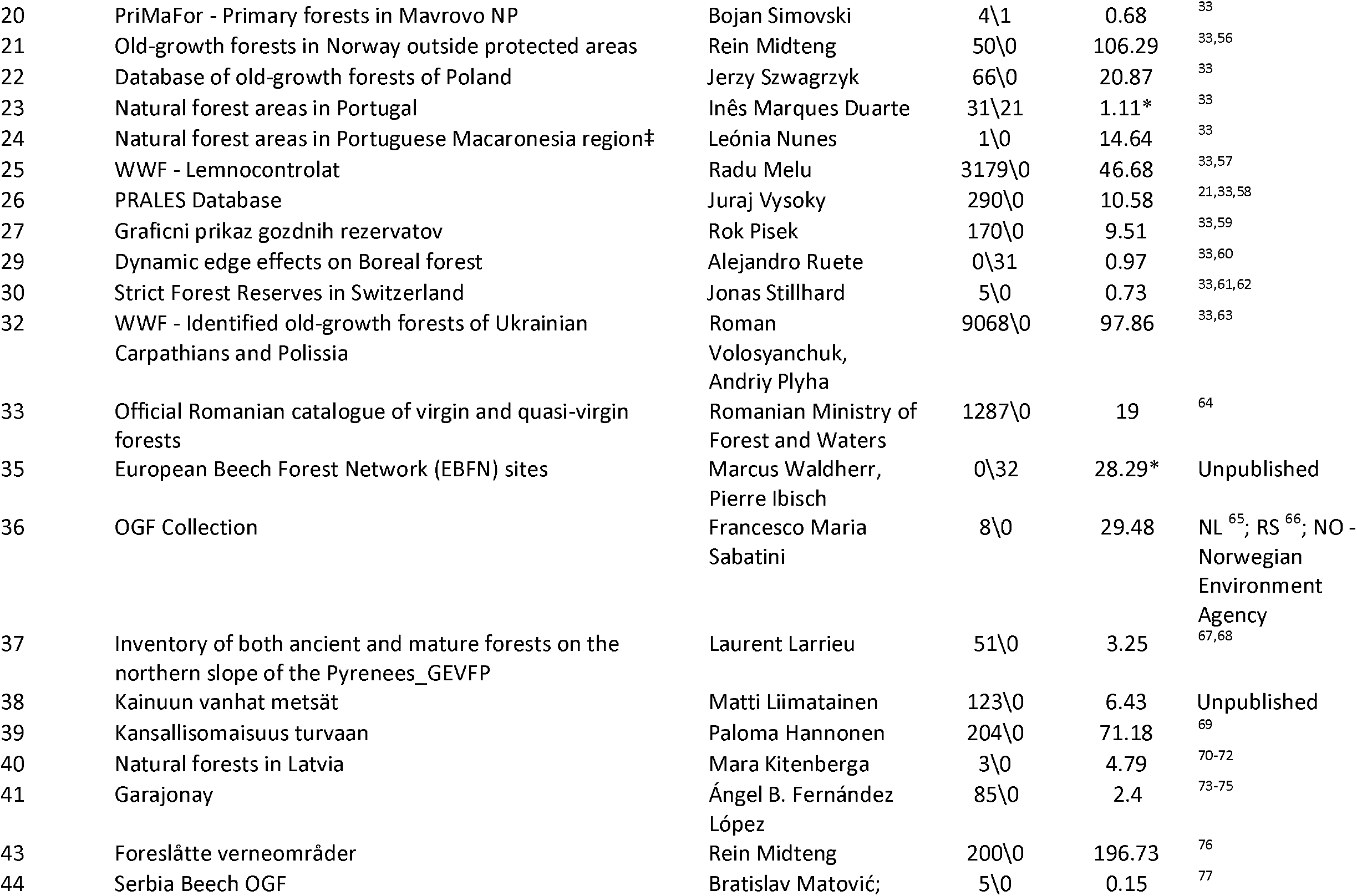
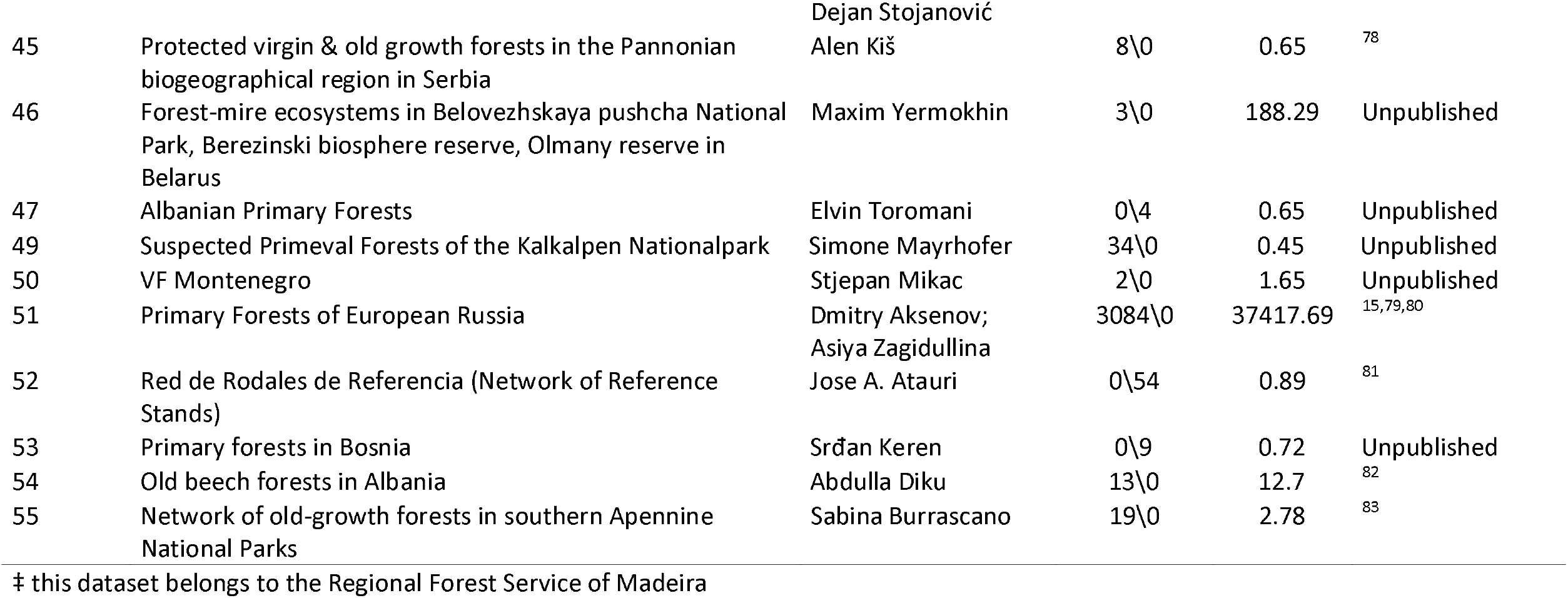
Synthetic description of datasets retrieved. ID codes are not consecutive. * Some point features have no information on forest patch area. † Overlapping areas across different datasets e-counted.

**Figure 2.**
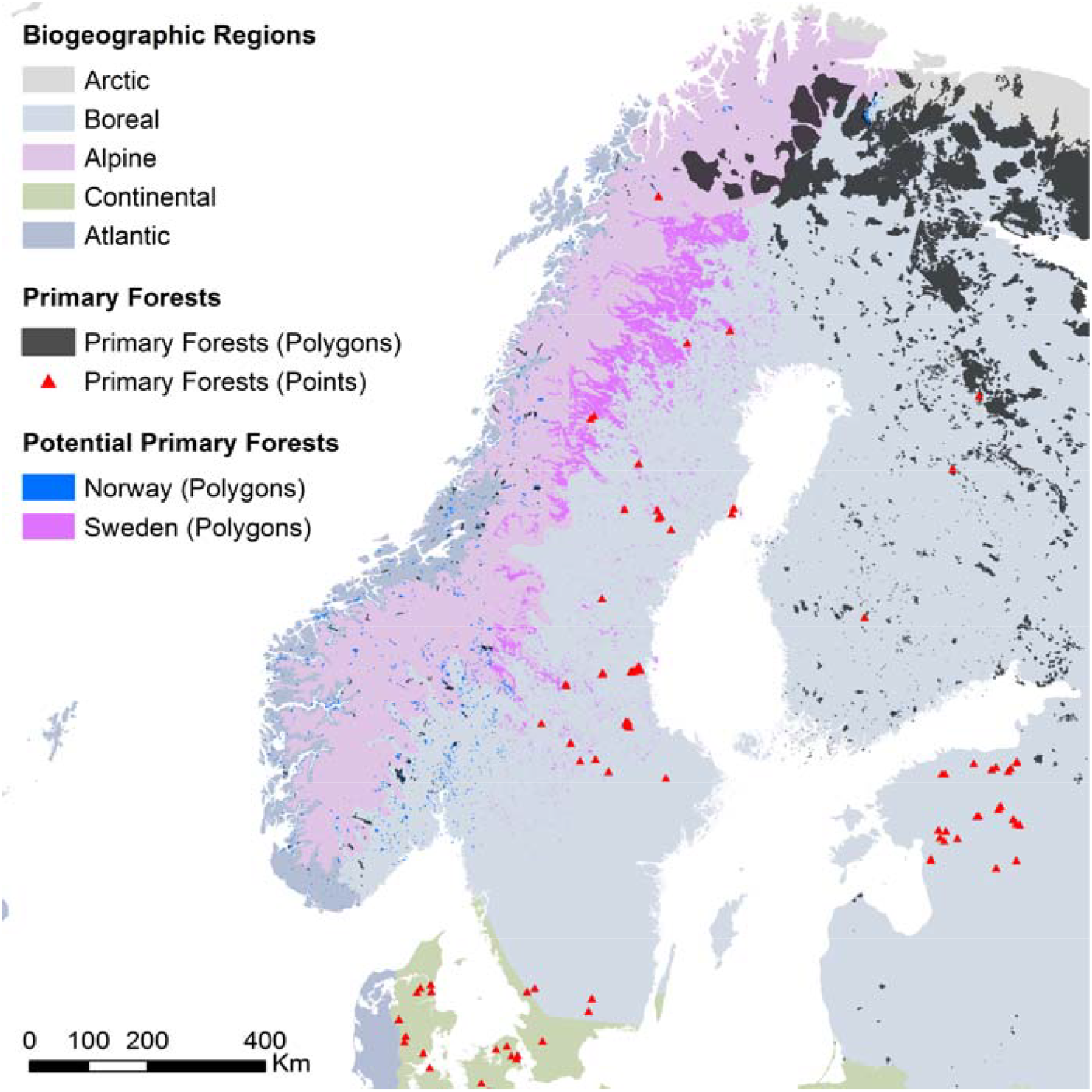
Overview of the maps of potential primary forests of Sweden and Norway.

## Methods

To define primary forests, we integrated the FAO definition of primary forests^1^, with the framework proposed by Buchwald [^35^]. In this framework, the term “primary forest” includes all forests where the signs of former human impacts, if any, are strongly blurred due to decades (at least 60-80 years) without forestry operations^35^. ‘Primary forests’ is therefore an umbrella term to include forests with different levels of naturalness, such as primeval, virgin, near-virgin, old-growth and long-untouched forests^35^. Our definition of primary forests, therefore, does not imply that these forests were never cleared or disturbed by humans, and includes, beside late-successional forests, also early seral stages and young forests that originated after natural disturbances and natural regeneration, without subsequent management. In case of large forest tracts (>250 ha) with high naturalness, our definition also allows forest polygons that include land temporarily or permanently not covered by trees.

To create the EPFD v2.0, we first expanded and updated the literature review on primary forests we had originally carried out for EPFD v1.0^33^, which only considered the period 2000-2017, and did not consider European Russia. Specifically, we added all scientific studies published between January 2000 and January 2019 for Russia, and those published in 2017-2019 for the rest of Europe. We identified relevant publications in the ISI Web of Knowledge using the search terms “(primary OR virgin OR old-growth OR primeval) AND forest*” in the title field. In line with [^33^], we deliberately excluded terms such as “unmanaged” (meaning: not under active management), “ancient” (never cleared for agriculture) or “natural” (stocked with naturally regenerated native trees). These terms indicate conditions that are necessary, but not sufficient for considering a forest as primary. Finally, we refined our search using geographical and subject filters. The literature search returned 122 candidate papers. After screening their content, we added 23 additional primary forest stands (10 in European Russia, 13 in the rest of Europe), from 13 studies (four from European Russia, and nine from the rest of Europe).

Building the EPFD v1.0^33^ involved reaching out to 134 forest experts. For v2.0 we contacted an additional 75 experts with knowledge on forests or forestry, and invited them to add spatially-explicit data on primary forests to our database. We focussed on experts from geographical regions poorly covered in v1.0. We received 56 answers, which led to the incorporation of 19 new datasets in our map. Given the context-dependency of definitions used in regional mapping projects, new datasets were only included if we could find an explicit equivalence between country-specific forest definitions and our definition framework^35^.

We integrated all data into a geodatabase, which contains primary forests either as polygons (if information on the forest boundary was available) or point locations (when having only a centroid). We set 0.5 hectares as minimum mapping unit. If available, we included a set of basic descriptors for each patch: name, location, naturalness level (based on [^35^]), extent, dominant tree species, disturbance history and protection status. In total, our map harmonized 48 regional-to-continental datasets of primary forests (Table 2). All data is open-access^34^. Besides, we retrieved three additional datasets that we kept confidential, either for conservation or copyright reasons. These datasets are: ‘Hungarian Forest Reserve monitoring’ (ID 17, custodian: Ferenc Horváth); ‘Ancient and Primeval Beech Forests of the Carpathians and Other Regions of Europe’^36,37^ (ID 34, custodian: UNESCO), and ‘Potential OGF and primary forest in Austria’ (ID 48, custodian: Matthias Schickhofer). Additional non-open access polygons also exist for the dataset ‘Strict Forest Reserves in Switzerland’ (ID 30, custodian: Jonas Stillhard). These data are referred to here for transparency, but are not included in the statistics and summaries reported in this paper.

### Post-Processing

To provide common descriptions for all features contained in the geodatabase, we integrated the basic descriptors detailed above with a range of attributes derived by intersecting all polygons or points with layers of: 1) biogeographical regions, 2) protected areas, 3) forest type, and 4) forest cover.

Overlaying the map of biogeographical region^84^ returned ten classes: 1. Alpine, 2. Arctic, 3. Atlantic, 4. Black Sea, 5. Boreal, 6. Continental, 7. Macaronesia, 8. Mediterranean, Pannonian, 10. Steppic. Information on protection status and time since onset of protection was based on the World Database of Protected Areas (WDPA)^85^. We simplified the original IUCN classification to three classes: 1. strictly protected – (IUCN category I); 2. protected – (IUCN categories II-VI + not classified); 3. not protected. We considered a primary forest patch as protected if >75% of its surface was within a WDPA polygon. When better information on the protection status of a forest patch was available directly from data contributors, we gave priority to this source. Forest type was based on the 14 forest categories defined by the European Environmental Agency^75^. The spatial information was derived by simplifying the map of Potential Vegetation types for Europe^86^, after creating a cross-link table^23^. The 13 categories comprise: 1. Boreal forest; 2. Hemiboreal forest and nemoral coniferous and mixed broadleaved፧coniferous forest; 3. Alpine coniferous forest; 4. Acidophilous oakwood and oak፧birch forest; 5. Mesophytic deciduous forest; 6. Lowland to submountainous beech forest; 7. Mountainous beech forest; 8. Thermophilous deciduous forest; 9. Broadleaved evergreen forest; 10. Coniferous forests of the Mediterranean, Anatolian and Macaronesian regions; 11. Mire and swamp forest; 12. Floodplain forest; 13. Non፧riverine alder, birch or aspen forest. For each primary forest patch, we reported the two most common forest categories. Finally, we extracted for each polygon the actual share covered by forest. We did this, because larger primary forest polygons in high naturalness classes can encompass land temporarily or permanently not covered by trees. We used a tree cover density map for the year 2010 for these regions from [^87^]. All post-processing was performed in R (v3.6.1)^88^.

### Data Gaps

To assess the completeness of our map, we calculated the ratio between the area of primary forest in our database at country level, and the estimated area of “forest undisturbed by man” from the indicator 4.3 in the Forest Europe report^89^. Although the definition of “forest undisturbed by man” in [^89^] is consistent with our definition of primary forest, it must be noted that these country-level estimates stem from national inventories or studies based on different interpretations, and the data quality varies from country to country. The comparison presented here should, therefore, be taken with caution (Figure 3).

**Figure 3.**
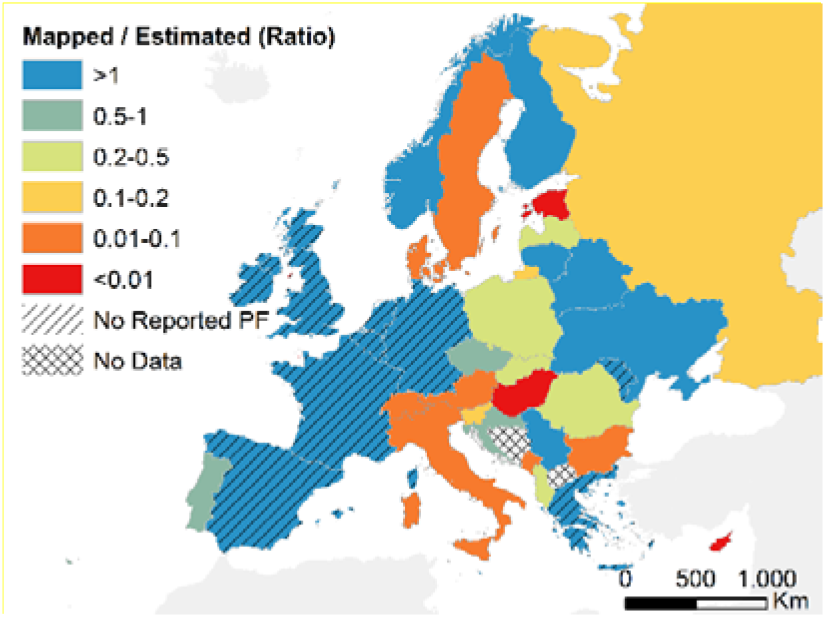
Estimation of data completeness. Ratio between the total primary forest area in the EPFD v2.0 and the^89^ country estimate of ‘forest undisturbed by man’ (indicator 4.3) from Forest Europe. Parallel hatching represents countries where Forest Europe reports either no forest undisturbed by man (‘No Reported PF’), or where data on forests undisturbed by man are missing (‘No Data’).

Forest Europe reports no primary forest for some western European countries (Spain, France, Belgium, Netherlands, Germany, United Kingdom and Ireland), although for most of these countries we did find information on at least a handful of primary forest sites. The coverage of our map was also higher than expected for some Eastern European countries (e.g., Ukraine, Belarus, Lituania), as well as Norway and Finland, known for hosting large areas of primary forests. Data completeness was lower for some central European countries. In the case of Czechia, Slovakia, Poland and Romania, our data only accounted for 20-100% of the country-level estimates from [^89^]. For Austria, Switzerland and Hungary, instead, data on primary forests exists but it is not currently open-access, and therefore not considered here. The largest data gaps were in Sweden, Italy, Bulgaria, Estonia, Denmark and Russia, where our map accounted for less than 10% of the primary forest reported in [^89^]. The low data completeness found for Denmark likely depends on the inclusion of minimum-intervention forest reserves in [^83^] that were harvested until then and therefore do not qualify as primary forests according to our definition.

### Potential Primary forests of Sweden and Norway

For Sweden and Norway, where abundant geographic information was available on forest distribution, we created maps of potential (yet unconfirmed) primary forests, as a way to complement our map. For Sweden, we derived a workflow to create a map of potential primary forests as detailed in Figure 4. This yielded 14,300 polygons covering a total area of 2.4 Mha.

**Figure 4.**
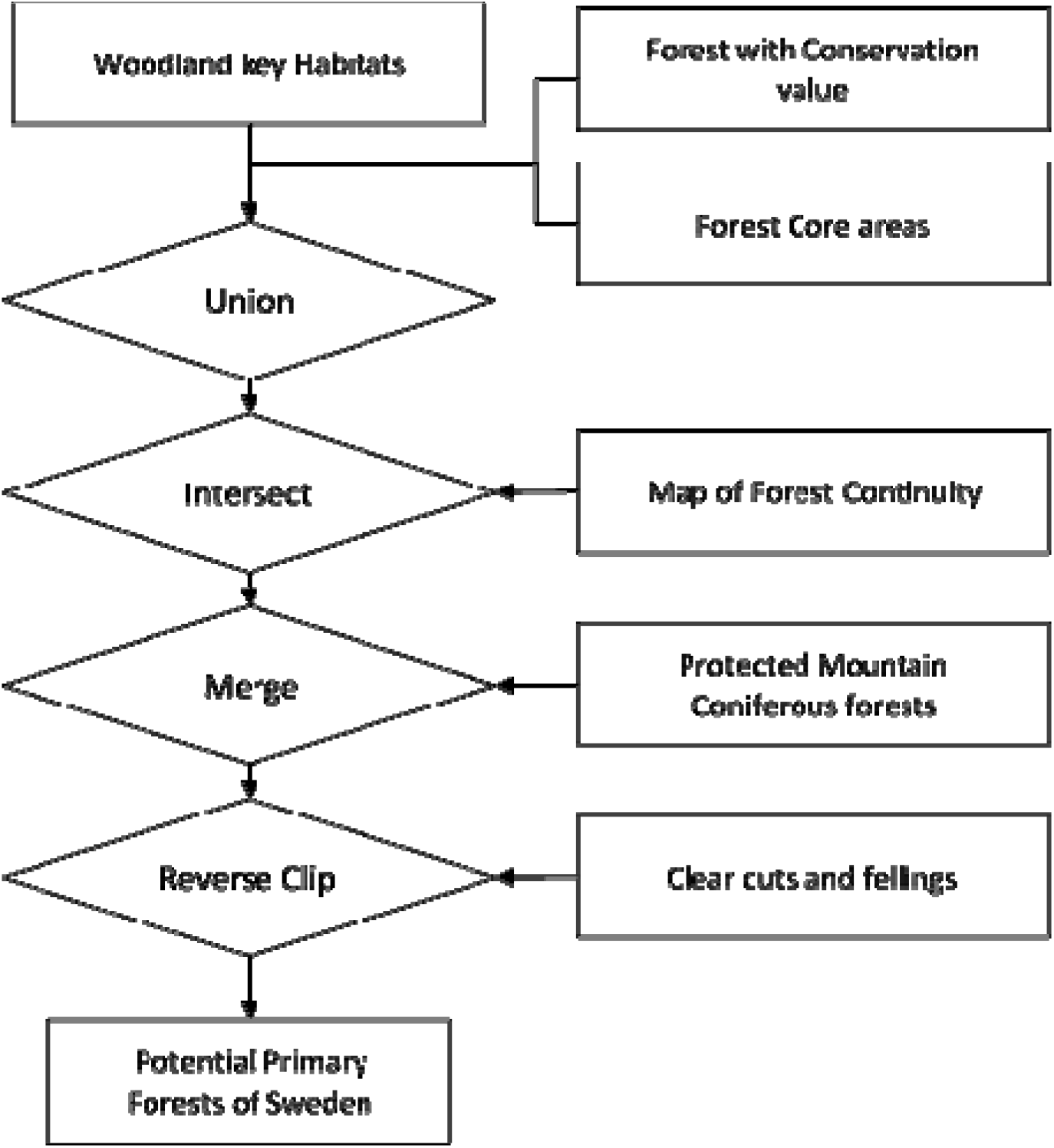
Workflow and data sources for the map of potential primary forests in Sweden. Data on woodland key habitats derive from [^90,91^]; forest with conservation value from [^92,93^], forest core areas from [^94^], continuity forests from [^95,96^], protected mountain coniferous forests from [^97^], clear cuts and fellings from [^98^].

For Norway, even though we were able to include two datasets of confirmed primary forests, additional primary forest is expected to exist. Therefore, we derived a map of potential primary forests, based on the “Viktige Naturtyper” dataset from the Norwegian Environment Agency^99^, which maps different habitat types of high conservation value both inside and outside forested areas. We extracted all polygons larger than 10 ha classified as “old forest types” (=“gammelskog”), i.e., forests that have never been clearcut and are in age classes of 120 years or older. This yielded 2,103 polygons covering a total area of 0.1 Mha.

### Data Records

The EPFD v2.0^34^ is composed of 48 individual datasets (Table 2), which we harmonized into two aggregated feature classes, after excluding all duplicated\overlapping polygons across individual datasets.

1. EU_PrimaryForests_Polygons_OA_v20

፧ Composite feature class combining the forest patches classified as “primary forest” based on polygon data sources described in Table 2
፧ Data type: Polygon Feature Class

1. EU_PrimaryForests_Points_OA_v20

፧ Composite feature class combining forest locations classified as “primary forest”, based on point data sources described in Table 2. Only points not overlapping with polygons in (1) reported.
፧ Data type: Point Feature Class

The individual datasets are also included in the geodatabase, inside the feature datasest ‘European_PrimaryForests’. The dataset is stored in Figshare (https://doi.org/10.6084/m9.figshare.13194095.v1)^34^. The file format is ESRI personal geodatabase (.mdb). Each feature class in the geodatabase follows the structure described in Table 3.

**Table 3.**
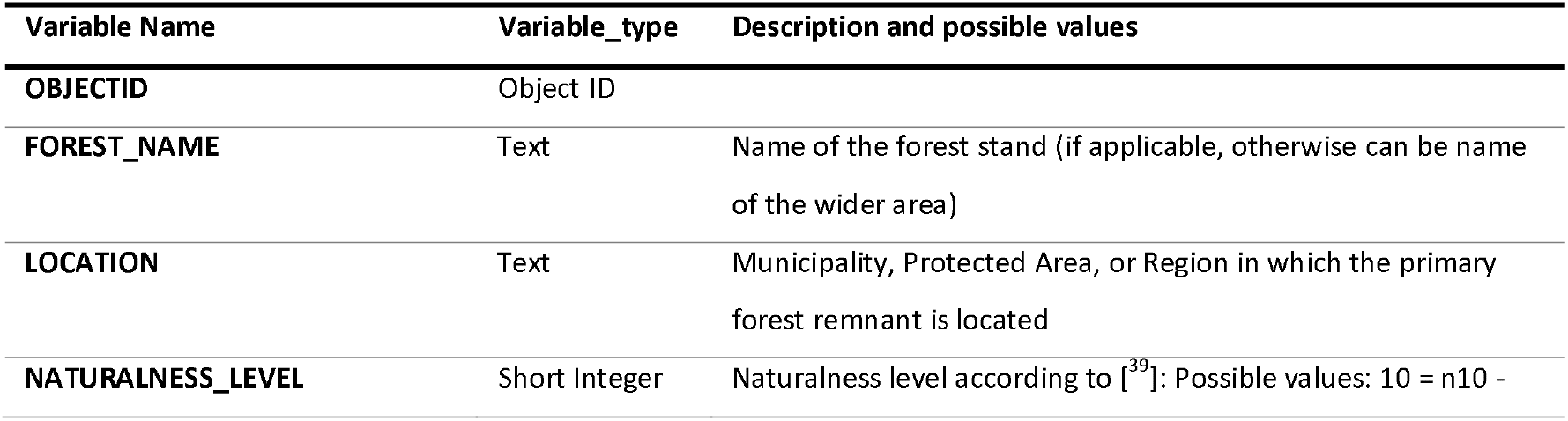
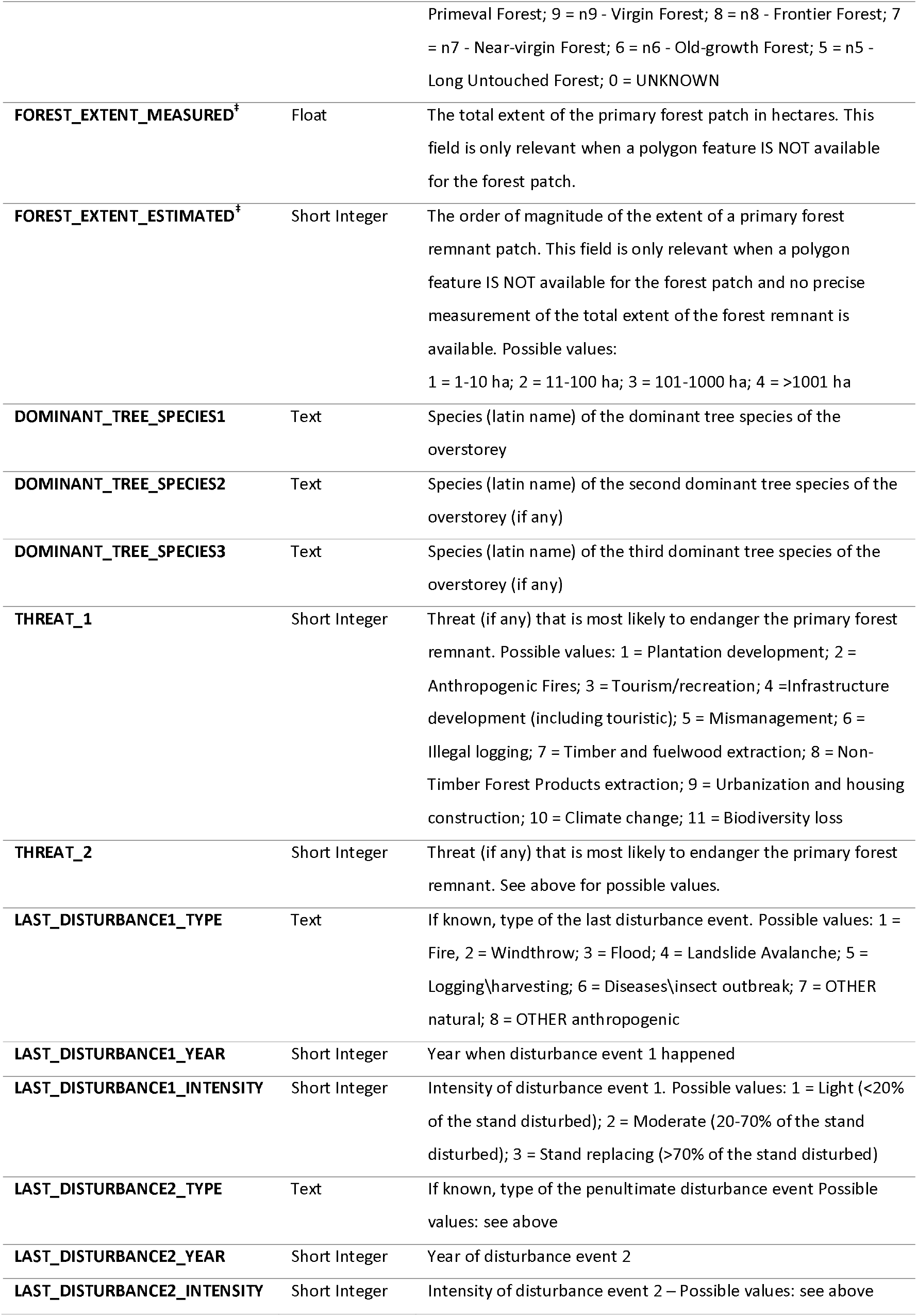
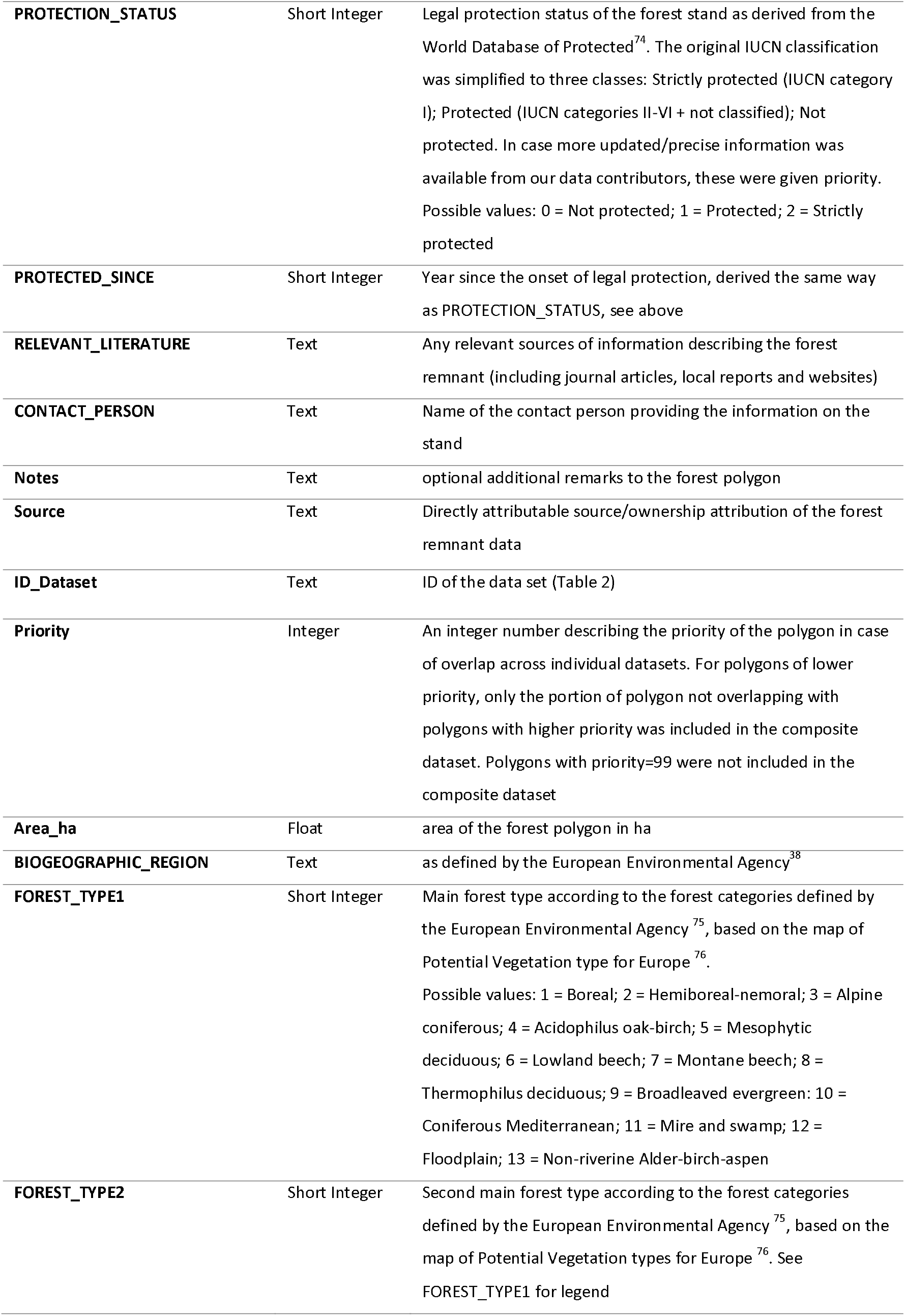
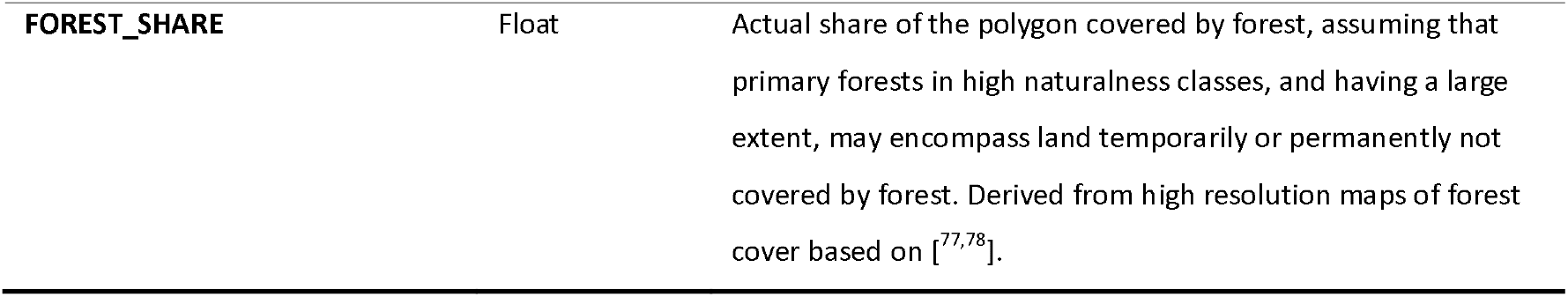
Spatial attributes of the feature classes of primary forests. ‡ - Only for point feature classes.

### Technical Validation

Although we had no direct control of the raw data contained in our database, the fact that all our information on primary forest locations derives either from peer-reviewed scientific literature, or were field-checked by trained researchers and/or professionals suggests high data reliability. We made sure to have a common understanding with data contributors about forest definitions [i.e., ^1,35^], and only included a dataset in the EPFD if we could find an explicit equivalence with the forest definitions we used.

To further assess data reliability, we carried out a robustness check using the open-access Landsat archive and the LandTrendr disturbance detection algorithm^100,101^, both implemented in Google Earth Engine^102^ (Figure 5). Specifically, we 1) quantified the proportion of polygons in our map, which underwent disturbance between 1985 and 2018, i.e., Landsat 5 operating time, 2) visually checked a subset of these disturbed polygons, to quantify the prevalence of anthropogenic vs. natural disturbance, and 3) extrapolated these results to the whole database to provide an estimation of the proportion of polygons in our map not meeting the necessary, but not sufficient, condition for being classified as primary (i.e. not being affected by anthropogenic disturbance within the last 35 years).

**Figure 5.**
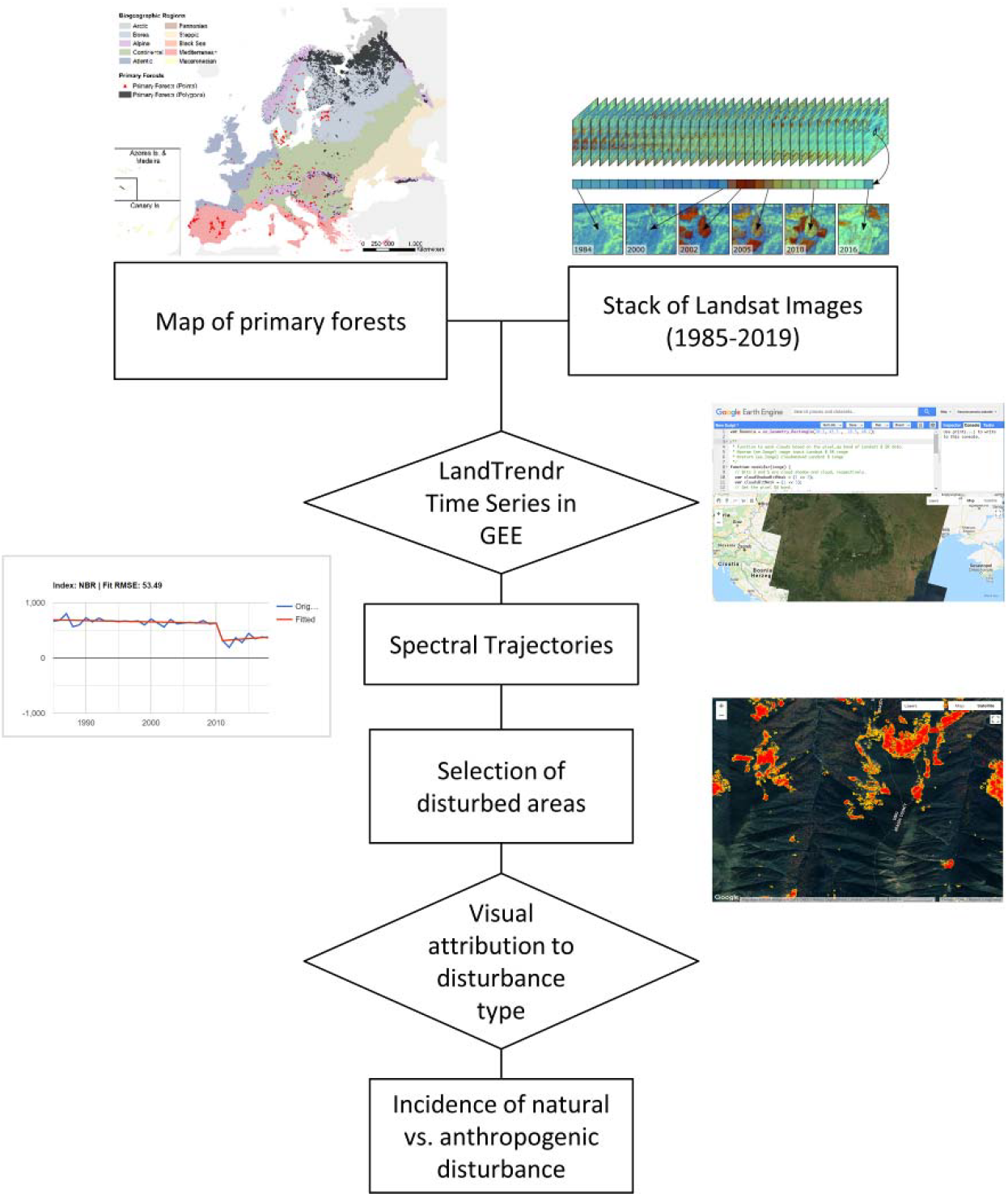
Workflow of the data robustness check.

For each polygon contained in the map of primary forests, we extracted the whole stack of available Landsat images (~1985-today), and ran the LandTrendr^103^ algorithm. LandTrendr identifies breakpoints in spectral time series, separates periods of disturbance or stability, and records the years in which disturbances occurred. To avoid problems due to cloud cover, changes in illumination, and atmospheric condition, we used all available images from the growing season of each year (1 May through 15 September) to derive yearly composite images^104^. As our spectral index, we used Tasseled Cap Wetness (TCW), as this index is particularly sensitive to forest structure^105^, is robust to spatial and temporal variations in canopy moisture^106^, and consistently outperforms other spectral indices, including Normalized Difference Vegetation Index^103^, for detecting forest disturbance^100,107,108^.

After running LandTrendr, we eliminated noise by applying a minimum disturbance threshold (2 ha). We then visually inspected a subset of primary forest polygons highlighted as ‘disturbed’ by LandTrendr. Based on the spectral and physical characteristics of the disturbed patch (brightness, shape, size), and on ancillary information derived from very-high-resolution images available in Google Earth, we assigned disturbance agents as either anthropogenic (i.e., forest harvest, infrastructure development) or natural (e.g., windstorm, bark beetle outbreak, fire; Figure 6, Figure 7).

**Figure 6.**
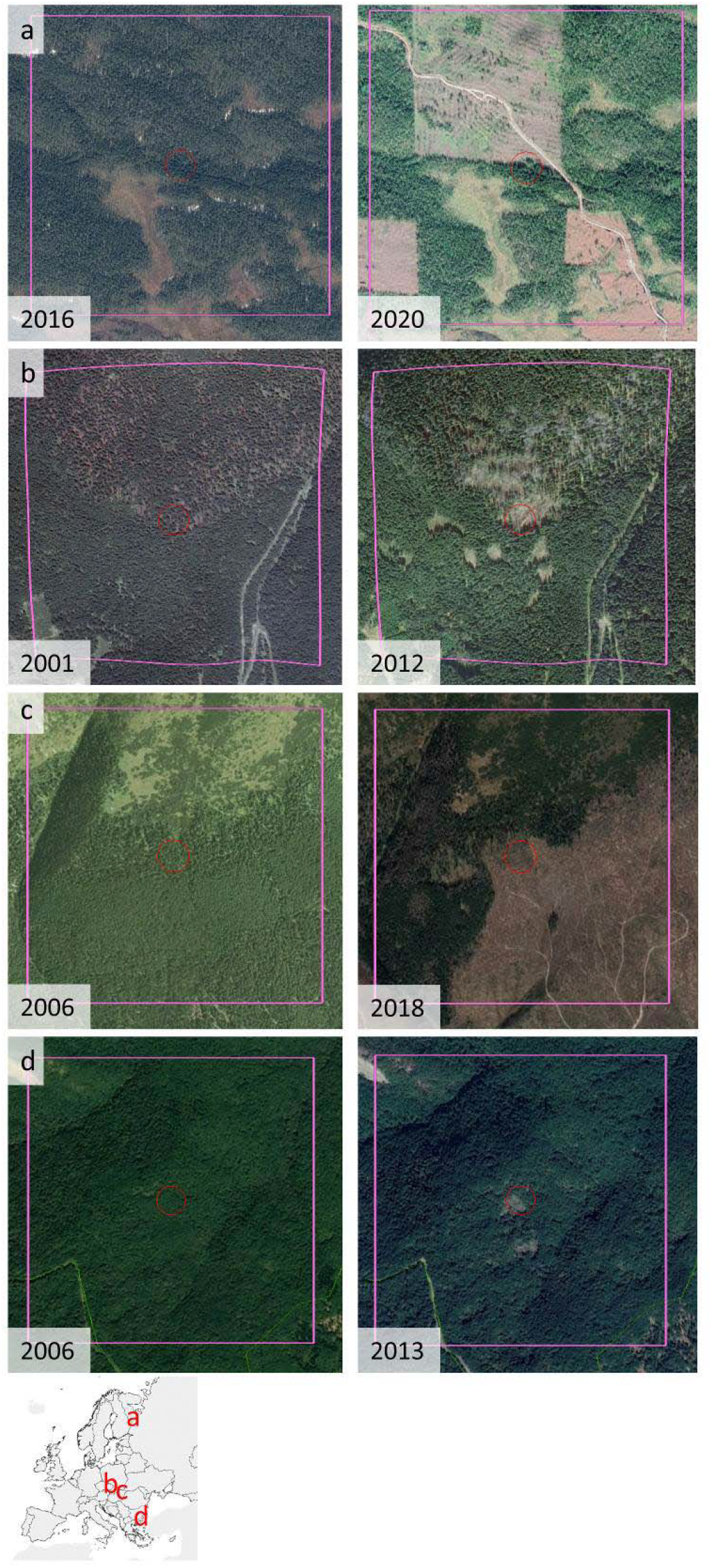
Examples of disturbed polygons, as detected by LandTrendr, before (left) and after (right) disturbance. a) clearcuts in the Russian Republic of Karelia; b) natural disturbance in Babia Gora, Slovakia; c) clear-cuts in Tatra National Park in Slovakia; d) natural disturbance in the southern Bourgas Province of Bulgaria. Red circles have a radius of 50 m; pink squares have a side of 1 km. Image credits: Google Earth.

**Figure 7.**
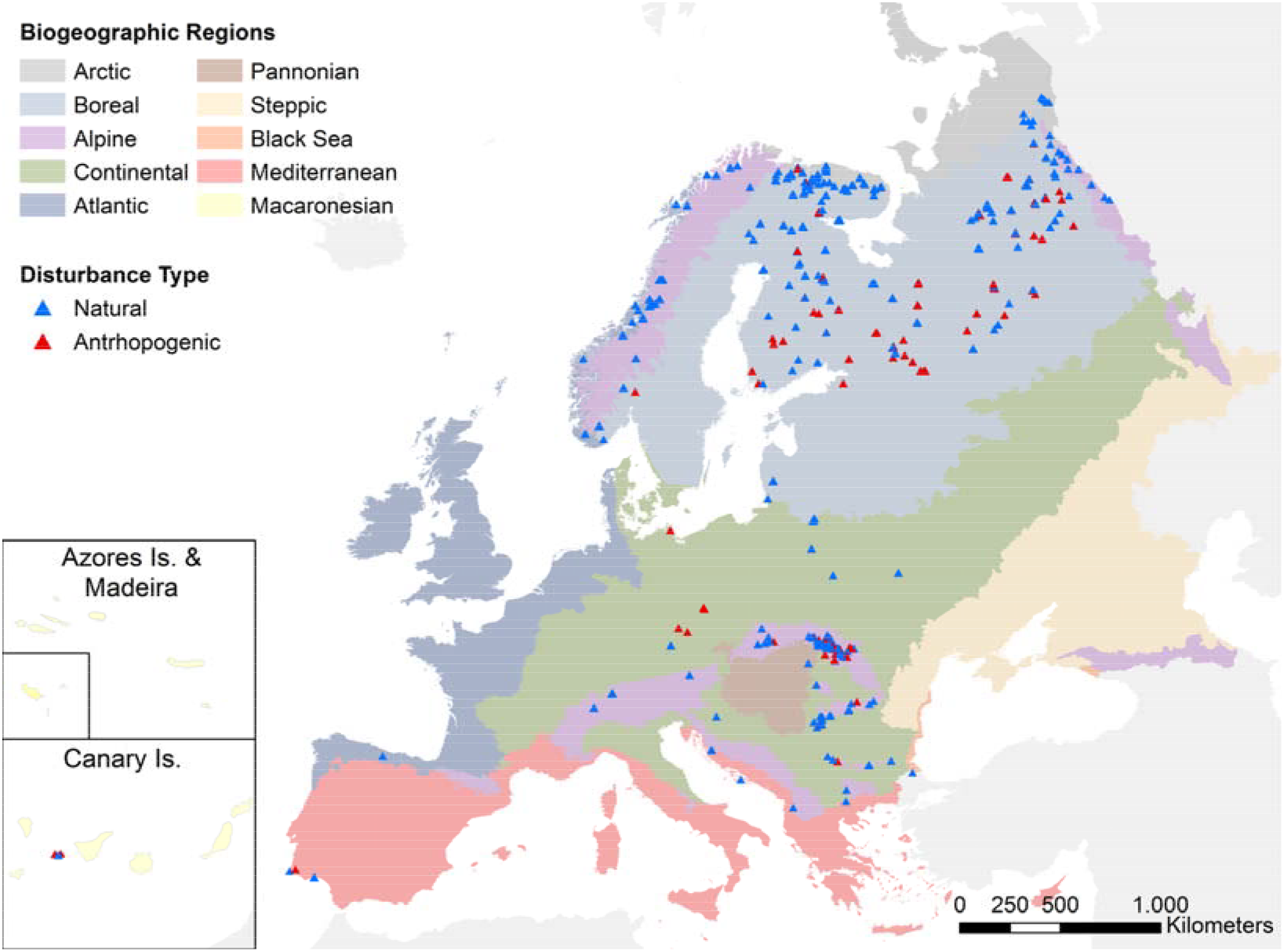
Geographical distribution of naturally vs. anthropogenically disturbed polygons, as resulting from a visual check of 712 disturbed polygons.

Out of the 17,309 polygons we checked, LandTrendr returned 4,734 polygons (27.3% of total) which experienced major disturbances between 1985 and 2018. The proportion of disturbed area was greater than 10% in 2,904 polygons. We visually checked 20% of the disturbed polygons in each biogeographic region, up to a maximum of 100 polygons. Depending on the size of the polygons, we inspected up to 5 pixels with a minimum distance of 1km. As a result, we visually inspected a total of 712 pixels across 268 primary forest polygons, therefore validating 1.5% of the total number of polygons and 5.7% of the disturbed polygons. We attributed a total of 149 pixels, across 61 primary forest polygons, to anthropogenic disturbance, (i.e., 22.7%, standard error = 2.5%) of the polygons we checked (Table 4, Figure 7). We thus estimated the total number of primary forest polygons being anthropogenically disturbed by multiplying the total number of polygons by the proportion of disturbed polygons (27.3%) and the share of these disturbed polygons attributed to anthropogenic causes (22.7%). This suggests our map contains 1,077 anthropogenically disturbed polygons (95% CIs [847, 1323]), which corresponds to 6.2% (95% CIs [4.9%, 7.6%]) of the total number of polygons. Disturbed polygons were concentrated in the Russian Federation (especially in Archangelsk region, Karelia and Komi republics), Southern Finland, and the Carpathians (Figure 7; Table 4). The Boreal and Alpine biogeographical regions had the highest number of disturbed polygons (both in total, and when considering only those with evident anthropogenic disturbance). The regions with the highest share of anthropogenically disturbed polygons was the Macaronesian, followed by the Continental and Boreal. Please note, that this robustness check should be considered as a low estimate, because only the disturbance events with a magnitude sufficient to be captured with LandTrendr and occurring in 1985-2018 could be identified.

**Table 4.** Results of robustness check, summarized by biogeographical region. † The number of disturbed polygons is higher than the number of polygons because some polygons expanding over more than one biogeographical region are double counted. PF – Primary Forest.

| Biogeographic region | Num. PF polygons (1) | Num. disturbed PF polygons (2) | Num. disturbed PF polygons checked (3) | Num. of (3) with evident anthropogenic disturbance (4) | Share of (3) anthropogenically disturbed (4/3) % |
| --- | --- | --- | --- | --- | --- |
| Alpine | 11,734 | 1,096 | 102 | 23 | 22.55 |
| Arctic | 96 | 105<sup>†</sup> | 20 | 0 | 0.00 |
| Atlantic | 83 | 48 | 13 | 0 | 0.00 |
| Black Sea | 19 | 6 | 1 | 0 | 0.00 |
| Boreal | 4,074 | 3,334 | 110 | 30 | 27.27 |
| Continental | 1,100 | 105 | 21 | 6 | 28.57 |
| Macaronesia | 27 | 8 | 2 | 1 | 50.00 |
| Mediterranean | 132 | 27 | 5 | 1 | 20.00 |
| Pannonian | 39 | 4 | 1 | 0 | 0.00 |
| Steppic | 5 | 1 | 0 | 0 | 0.00 |

### Usage Notes

All data files are referenced in a geographic coordinate system (lat/long, WGS 84 - EPSG code: 4326). The provided files are in a personal geodatabase, and can be accessed and displayed using standard GIS software such as: QGIS (www.qgis.org/en).

All datasets listed in Table 2 are freely available in Figshare (https://doi.org/10.6084/m9.figshare.13194095.v1)^34^ with a Creative Commons CC BY 4.0 license. Three additional non open-access datasets are available on request after approval of the respective copyright holders. These datasets are: ‘Hungarian Forest Reserve monitoring’ (ID 17, custodian: Ferenc Horváth); ‘Ancient and Primeval Beech Forests of the Carpathians and Other Regions of Europe’^36,37^ (ID 34, Custodian: UNESCO), and ‘Potential OGF and primary forest in Austria’ (ID 48, custodian: Matthias Schickhofer). The same conditions apply for additional data from the dataset ‘Strict Forest Reserves in Switzerland’ (ID 30, custodian: Jonas Stillhard). Comments and requests of updates for the dataset are collected and discussed in the GitHub forum: https://github.com/fmsabatini/PrimaryForestEurope.

## Code Availability

The code to reproduce the post-processing is available in Figshare

(https://doi.org/10.6084/m9.figshare.13194095.v1)^34^. The dataset contains five scripts:

- 00_ComposeMap.R – Identifies overlapping polygons across individual datasets
- 01_CreateComposite_Points.py – creates the composite point feature class.
- 02_CreateComposite_Polygons.py – creates the composite polygon feature class.
- 03_PostProcessing.R – Extract additional information on each primary forest
- 04_Add_Postprocessing.py – Imports post-processing output into the geodatabase
- 05_Summary_stats.R – Calculates summary statistics of primary forests

Python (.py) scripts were run in ESRI ArcGIS (v10.5) and are available also as ArcGIS Models inside the Geodatabase. The remaining (.R) scripts were run using R (v4.0).

## Acknowledgements

This project was funded by Frankfurt Zoological Society (FZS; project ETN-WIE-FZS-001) and the European Commission (Marie Sklodowska-Curie fellowship to FMS, project FORESTS & CO, #658876). The Italian dataset was supported by funding from the Department for Nature Protection of the Italian Ministry of the Environment, Land and Sea Protection. The GEVFP's French dataset was supported by funding from EU, RF and 'Conseil Régional Midi-Pyrénées’. MM gratefully acknowledges the institutional project EVA No.CZ.02.1.01/ 0.0/0.0/16_019/0000803 and MŠMT project LTT20016. This work would not have been possible without all those who responded to the questionnaires and those who collected all the data presented here. A special thanks goes to all data contributors, especially those who are not coauthors of this paper (in alphabetic order): Paloma Hannonen (Dataset 39), Matti Liimatainen (Dataset 38), Simone Mayrhofer (Dataset 49), Daniel Vallauri (Dataset 13) and Juraj Vysoky (Dataset 26). We also thank Teresa De Marzo for support during the validation process.

## Author contributions

The original idea for the database is from FMS, TK and ZK. FMS and HB harmonized the datasets, and conducted the literature review. DA, JAA, EB, SB, EC, AD, IMD, ABF, MG, NG, FH, SK, MK, AKi, AKr, PLI, LL, FL, BM, RNM, PM, SM, RM, MM, GM, MP, RP, LN, AR, MS, BS, JS, DS, JS, O-PT, ET, RV, TV, MW, MY, TZ, AZ contributed data. FMS, HB, and TK created the first draft of the manuscript and all co-authors contributed substantially to its revision.

## Competing interests

The authors declare no competing financial interests.

## Notes

https://doi.org/10.6084/m9.figshare.13194095.v1

### Competing Interest Statement

The authors have declared no competing interest.

### Summary of Updates

Version submitted to Scientific Data after first revision. In this version, only open-access data sets are described and summarized, while confidential data sets are only mentioned.

## References

1 FAO. Global Forest Resources Assessment 2015. Terms and definitions. (FAO, 2015).

2 Lansstyrelsen. Formellt skyddad skogi Norrbottens län. Report No. 0283–9636, (2018).

3 Watson, J. E. M. et al. The exceptional value of intact forest ecosystems. Nat. Eco. Evo., 1 (2018).

4 European Commission. in COM(2020) 380 final (Brussels, 2020).

5 Vandekerkhove, K. et al. Reappearance of Old-Growth Elements in Lowland Woodlands in Northern Belgium: Do the Associated Species Follow? Silva Fenn. 45, 909–935, doi:10.14214/sf.78 (2011).

6 Di Marco, M., Ferrier, S., Harwood, T. D., Hoskins, A. J. & Watson, J. E. Wilderness areas halve the extinction risk of terrestrial biodiversity. Nature, 1–4 (2019).

7 Frey, S. J. K. et al. Spatial models reveal the microclimatic buffering capacity of old-growth forests. Science Advances 2, e1501392, doi:10.1126/sciadv.1501392 (2016).

8 Zhou, G. Y. et al. Old-growth forests can accumulate carbon in soils. Science 314, 1417–1417, doi:10.1126/science.1130168 (2006).

9 Burrascano, S., Keeton, W. S., Sabatini, F. M. & Blasi, C. Commonality and variability in the structural attributes of moist temperate old-growth forests: A global review. For. Ecol. Manag. 291, 458–479, doi:http://dx.doi.org/10.1016/j.foreco.2012.11.020 (2013).

10 Bauhus, J., Puettmann, K. & Messier, C. Silviculture for old-growth attributes. For. Ecol. Manag. 258, 525–537, doi:10.1016/j.foreco.2009.01.053 (2009).

11 Moore, K. D. In the shadow of the cedars: the spiritual values of old-growth forests. Conserv. Biol. 21, 1120–1123 (2007).

12 FOREST EUROPE. State of Europe’s Forests 2015. (Ministerial Conference on the Protection of Forests in Europe, Madrid, 2015).

13 Ceccherini, G. et al. Abrupt increase in harvested forest area over Europe after 2015. Nature 583, 72–77, doi:10.1038/s41586-020-2438-y (2020).

14 Levers, C. et al. Drivers of forest harvesting intensity patterns in Europe. For. Ecol. Manag. 315, 160–172 (2014).

15 Potapov, P. et al. The last frontiers of wilderness: Tracking loss of intact forest landscapes from 2000 to 2013. Science Advances 3, doi:10.1126/sciadv.1600821 (2017).

16 Schickhofer, M. & Schwarz, U. Inventory of Potential Primary and Old-Growth Forest Areas in Romania (PRIMOFARO). Identifying the largest intact forests in the temperate zone of the European Union. (Euronatur Foundation, 2019).

17 Knorn, J. et al. Continued loss of temperate old-growth forests in the Romanian Carpathians despite an increasing protected area network. Environ. Conserv. 40, 182–193 (2013).

18 Court of Justice of the European Union. C-441/17 - Commission v Poland (Forêt de Białowieża) Judgment of the Court (Grand Chamber) of 17 April 2018 (2018).

19 Chylarecki, P. & Selva, N. Ancient forest: spare it from clearance. Nature 530, 419–419 (2016).

20 Earthsight. Complicit in corruption. How billion-dollar firms and EU governments are failing Ukraine's forests. (Earthsight, 2018).

21 Mikoláš, M. et al. Primary forest distribution and representation in a Central European landscape: Results of a large-scale field-based census. For. Ecol. Manag. 449, 117466, doi:https://doi.org/10.1016/j.foreco.2019.117466 (2019).

22 Hance, J. & But, S. s. IKEA Logging Old-growth Forest for Low-price Furniture in Russia. (2012). https://news.mongabay.com/2012/05/ikea-logging-old-growth-forest-for-low-price-furniture-in-russia/.

23 Sabatini, F. M. et al. Protection gaps and restoration opportunities for primary forests in Europe. Divers. Distrib. n/a, doi:10.1111/ddi.13158 (2020).

24 Adam, D. & Vrška, T. in Landscape Atlas of the Czech Republic (eds T Hrnčiarová, P Mackovčin, & I Zvara) 209 (Ministry of Environment and Silva Tarouca Research Institute, Prague–Silva Tarouca Research Institute for Landscape and Ornamental Gardening, 2009).

25 Bernadzki, E., Bolibok, L., Brzeziecki, B., Za̧jaczkowski, J. & Zybura, H. Compositional dynamics of natural forests in the Bialowieza National Park, northeastern Poland. J. Veg. Sci. 9, 229–238 (1998).

26 Hobi, M. L., Commarmot, B. & Bugmann, H. Pattern and process in the largest primeval beech forest of Europe (Ukrainian Carpathians). J. Veg. Sci. 26, 323–336 (2015).

27 Kral, K. et al. Local variability of stand structural features in beech dominated natural forests of Central Europe: Implications for sampling. For. Ecol. Manag. 260, 2196–2203, doi:10.1016/j.foreco.2010.09.020 (2010).

28 Veen, P. et al. Virgin forests in Romania and Bulgaria: results of two national inventory projects and their implications for protection. Biodivers. Conserv. 19, 1805–1819, doi:10.1007/s10531-010-9804-2 (2010).

29 Ibisch, P. L. & Ursu, A. (Greenpeace CEE Romania; Centre for Econics and Ecosystem Management, Eberswalde University for Sustainable Development; Geography Department, A. I. Cuza University of Iaşi, 2017).

30 Spracklen, B. D. & Spracklen, D. V. Identifying European Old-Growth Forests using Remote Sensing: A Study in the Ukrainian Carpathians. Forests 10, 127 (2019).

31 Diaci, J. in Proceedings of the invited lecturers’ reports presented at the COST E4 management committee and working groups meeting in Ljubljana, Slovenia. (Department of Forestry and Renewable Forest Resources Biotechnical Faculty).

32 Frank, G. et al. COST Action E27. Protected Forest Areas in Europe-analysis and harmonisation (PROFOR): results, conclusions and recommendations. (Federal Research and Training Centre for Forests, Natural Hazards and Landscape (BFW), 2007).

33 Sabatini, F. M. et al. Where are Europe’s last primary forests? Divers. Distrib. 24, 1426–1439, doi:10.1111/ddi.12778 (2018).

34 Sabatini, F. M. et al. European Primary Forest Database. figshare, https://doi.org/10.6084/m9.figshare.13194095.v1 (2020).

35 Buchwald, E. in Proceedings: Third expert meeting on harmonizing forest-related definitions for use by various stakeholders (Food and Agriculture Organization of the United Nations, 2005).

36 Britz, H. et al. Nomination of the “Ancient Beech Forests of Germany” as Extension to the World Natural heritage “Primeval Beech Forests of the Carpathians”. Nationale Naturlandschaften, Federal Republic of Germany. Nieden-stein: Specialised editing Cognitio Kommunikation & Planung (2009).

37 UNEP-WCMC & IUCN. Protected Planet: Ancient and Primeval Beech Forests of the Carpathians and Other Regions of Europe in Albania, Austria, Belgium, Bulgaria, Croatia, Germany, Italy, Romania, Slovakia, Slovenia, Spain and Ukraine, The World Database on Protected Areas (WDPA)/The Global Database on Protected Areas Management Effectiveness (GD-PAME), https://www.protectedplanet.net/903141 (2019).

38 Trotsiuk, V. et al. A mixed severity disturbance regime in the primary Picea abies (L.) Karst. forests of the Ukrainian Carpathians. For. Ecol. Manag. 334, 144–153, doi:http://dx.doi.org/10.1016/j.foreco.2014.09.005 (2014).

39 Kozák, D. et al. Profile of tree-related microhabitats in European primary beech-dominated forests. For. Ecol. Manag. 429, 363–374 (2018).

40 Svoboda, M. et al. Landscape-level variability in historical disturbance in primary Picea abies mountain forests of the Eastern Carpathians, Romania. J. Veg. Sci. 25, 386–401, doi:10.1111/jvs.12109 (2014).

41 Garbarino, M. et al. Gap disturbances and regeneration patterns in a Bosnian old-growth forest: a multispectral remote sensing and ground-based approach. Ann. For. Sci. 69, 617–625, doi:10.1007/s13595-011-0177-9 (2012).

42 Keren, S. et al. Comparative Structural Dynamics of the Janj Mixed Old-Growth Mountain Forest in Bosnia and Herzegovina: Are Conifers in a Long-Term Decline? Forests 5, 1243–1266 (2014).

43 Motta, R. et al. Structure, spatio-temporal dynamics and disturbance regime of the mixed beech–silver fir–Norway spruce old-growth forest of Biogradska Gora (Montenegro). Plant Biosyst. 149, 966–975, doi:10.1080/11263504.2014.945978 (2015).

44 Motta, R. et al. Development of old-growth characteristics in uneven-aged forests of the Italian Alps. Eur. J. For. Res. 134, 19–31, doi:10.1007/s10342-014-0830-6 (2015).

45 WWF Bulgaria. Forests in Bulgaria - Old growth forests, https://gis.wwf.bg/mobilz/en/#.

46 Panayotov, M. et al. Mountain coniferous forests in Bulgaria – structure and natural dynamics. (University of Forestry and Geosoft, 2016).

47 Blue Cat Team. Czech Natural Forests Databank, https://www.naturalforests.cz/czech-natural-forests-databank.

48 Lõhmus, A. & Kraut, A. Stand structure of hemiboreal old-growth forests: Characteristic features, variation among site types, and a comparison with FSC - certified mature stands in Estonia. For. Ecol. Manag. 260, 155–165, doi:http://dx.doi.org/10.1016/j.foreco.2010.04.018 (2010).

49 EEA. Developing a forest naturalness indicator for Europe. Concept and methodology for a high nature value (HNV) forest indicator. (EEA Technical report No 13/2014, Luxembourg: Publications Office of the European Union, 2014).

50 SYKE. Finnish Environment Institute, http://www.syke.fi/en-US/Open_information.

51 Rossi, M., Bardin, P., Cateau, E. & Vallauri, D. Forêts anciennes de Méditerrané e et des montagnes limitrophes: références pour la naturalité régionale. WWF France, Marseille, France, 144 (2013).

52 WWF France. Hauts lieux de naturalité en France, http://www.foretsanciennes.fr/proteger-mieux/france/.

53 Cateau, E. et al. Le patrimoine forestier des réserves naturelles. Focus sur les forêts à caractère naturel. Vol. cahier rnf 7 (Réserves Naturelles de France, 2017).

54 Bundesanstalt für Landwirtschaft und Ernährung. Datenbank Naturwaldreservate in Deutschland, https://www.naturwaelder.de/index.php?tpl=home.

55 Blasi, C., Burrascano, S., Maturani, A. & Sabatini, F. M. Old-growth forests in Italy. (Palombi Editori, 2010).

56 Myhre, T. Skogkur 2020. redningsplan for Norges unike skoger. WWF Verdens villmarksfond, Norges naturvernforbund, SABIMA (2012).

57 WWF Romania. Harta ariilor protejate din Romania, http://www.lemncontrolat.ro/en/home/.

58 Pralesy.Sk. Mapovanie pralesov Slovenska, http://en.pralesy.sk/.

59 Slovenia Forest Service. Gozdni rezervati, http://www.zgs.si/gozdovi_slovenije/o_gozdovih_slovenije/gozdni_rezervati/index.html.

60 Ruete, A., Snäll, T. & Jönsson, M. Dynamic anthropogenic edge effects on the distribution and diversity of fungi in fragmented old-growth forests. Ecol. Appl. 26, 1475–1485, doi:10.1890/15-1271 (2016).

61 Heiri, C., Wolf, A., Rohrer, L., Brang, P. & Bugmann, H. Successional pathways in Swiss mountain forest reserves. Eur. J. For. Res. 131, 503–518 (2012).

62 Brang, P., Heiri, C. & Bugmann, H. Waldreservate: 50 Jahre natürliche Waldentwicklung in der Schweiz. (Haupt, 2011).

63 WWF Ukraine. Virgin, Old-Growth and Natural Forests of Ukraine, http://gis-wwf.com.ua/.

64 Ministerului Apelor şi Pădurilor. Catalogul pădurilor virgine şi cvasivirgine din România, http://apepaduri.gov.ro/paduri-virgine/.

65 Alterra. Forest reserves, https://www.wur.nl/en/Research-Results/Projects-and-programmes/Humus-forms/Publications-Humus-forms/Forest-reserves.htm (2000).

66 Pantić, D. et al. Structural, production and dynamic characteristics of the strict forest reserve'Račanska šljivovica'on Mt. Tara. Glasnik Šumarskog fakulteta, 93–114 (2011).

67 Savoie, J. M. et al. Vieilles forêts pyrénéennes de Midi-Pyrénées. Deuxième phase. Evaluation et cartographie des sites. Recommandations. Rapport final. (Ecole d’Ingénieurs de PURPAN/DREAL Midi-Pyrénées, 2015).

68 Savoie, J. M. et al. Forêts pyrénéennes anciennes de Midi-Pyrénées. Rapport d’Etude de projet FEDER 2008–2011. 320 (Ecole d’Ingénieurs de PURPAN/DREAL Midi-Pyrénées, 2011).

69 WWF Finland. Kansallisomaisuus turvaan - valtion omistamia suojelun arvoisia metsä-ja suoalueita., (WWF Suomen raportteja, 2012).

70 Kitenberga, M. et al. A mixture of human and climatic effects shapes the 250-year long fire history of a semi-natural pine dominated landscape of Northern Latvia. For. Ecol. Manag. 441, 192–201 (2019).

71 Baders, E., Senhofa, S., Purina, L. & Jansons, A. Natural succession of Norway spruce stands in hemiboreal forests: case study in Slitere national park, Latvia. Baltic Forestry 23, 522–528 (2017).

72 Kokarēviča, I. et al. Vegetation changes in boreo–nemoral forest stands depending on soil factors and past land use during an 80 year period of no human impact. Can. J. For. Res. 46, 376–386 (2016).

73 Fernandez López, A. B. Parque Nacional de Garajonay, Patrimonio Mundial. (Organismo Autonomo Parques Nacionales, 2009).

74 TRAGSATEC. Segundo inventario ecológico del Parque Nacional de Garajonay. (Parque Nacional de Garajonay, 2006).

75 Fernández, A. B. & Gómez, L. Qué son los bosques antiguos de laurisilva. Su valor y situación en Canarias. La Gomera, entre bosques y taparuchas, 177–236 (2016).

76 BIO FOKUS. Narin lokalitetsdatabase, https://biofokus.no/narin/.

77 Matović, B. et al. Comparison of stand structure in managed and virgin european beech forests in Serbia. Šumarski list 142, 47–57 (2018).

78 Kiš, A., Stojšić, V., & Dinić, A. in 2nd International Symposium on Nature Conservation. Proceedings 373–382 (Institute for Nature Conservation of Vojvodina Province, Novi Sad, 2016).

79 Potapov, P. et al. Mapping the world’s intact forest landscapes by remote sensing. Ecol. Soc. 13 (2008).

80 Kobyakov, K. & Jakolev, J. Atlas of high conservation value areas, and analysis of gaps and representativeness of the protected area network in northwest Russia. Finnish Environment Institute (2013).

81 Europarc Spain. Mapa de rodales de referencia RedBosques, www.redbosques.eu ( 82 Diku, A. & Shuka, L. Pyjet e vjetër të ahut në shqipëri (Old Beech forests in Albania). (PSEDA ILIRIA http://iliria-al.org/publications/, 2017).

83 Burrascano, S. et al. It's a long way to the top: Plant species diversity in the transition from managed to old-growth forests. J. Veg. Sci. 29, 98–109, doi:10.1111/jvs.12588 (2018).

84 EEA. (ed Directorate-General for Environment (DG ENV) Council of Europe (CoE)) (2016).

85 UNEP-WCMC & IUCN. Protected Planet: The World Database on Protected Areas (WDPA) www.protectedplanet.net (2019).

86 Bohn, U. et al. Map of the natural vegetation of Europe. Explanatory text with CD-ROM., (German Federal Agency for Nature Conservation, Bonn, Germany, 2003).

87 Hansen, M. C. et al. High-resolution global maps of 21st-century forest cover change. Science 342, 850–853, doi:10.1126/science.1244693 (2013).

88 R: A language and environment for statistical computing v. 3.6.1 (R Foundation for Statistical Computing, Vienna, Austria., 2019).

89 FOREST EUROPE. Quantitative Indicators Country reports 2015, https://foresteurope.org/state-europes-forests2015-report/#1476295965372-d3bb1dd0-e9a0 (2015).

90 Skogsstyrelsen. Key Woodland Habitats, https://www.skogsstyrelsen.se/ (2003).

91 Frank, A. Inventering av nyckelbiotoper: resultat till och med 2003. (Skogsstyr., 2004).

92 Länsstyrelsen Västerbotten. LstAC Skogar med höga naturvärden ovan gränsen för fjällnära skog 2003–2015, https://ext-geodatakatalog.lansstyrelsen.se/GeodataKatalogen/ (2019).

93 Naturvårdsverket. Skyddsvärda statliga skogar, http://mdp.vicmetria.nu/miljodataportalen/GetMetaDataById?UUID=3919E66E-2E09-440D-9171-B5074DF0C0ED (2017).

94 Naturvårdsverket. Skogliga värdekärnor, http://gpt.vicmetria.nu/data/land/skogliga_vardekarnor_2016.zip (2016).

95 Naturvårdsverket. Preciserad kartering av kontinuitetsskog i Jämtlands län, http://gpt.vic-metria.nu/data/land/Preciserad_kskog_jamtland.zip (2019).

96 Ahlkrona, E., Giljam, C. & Wennberg, S. Kartering av kontinuitetsskogi boreal region. Metria AB på uppdrag av Naturvårdsverket., (2017).

97 Naturvårdsverket. Skyddad fjallbarrskog, https://gpt.vic-metria.nu/data/land/NMD/Skyddad_Fjallbarrskog.zip (2019).

98 Skogsstyrelsen. Utförda avverkningar, https://www.skogsstyrelsen.se/ (2014).

99 Miljødirektoratet. (2016).

100 Cohen, W. B., Yang, Z., Healey, S. P., Kennedy, R. E. & Gorelick, N. A LandTrendr multispectral ensemble for forest disturbance detection. Remote Sens. Environ. 205, 131–140 (2018).

101 Kennedy, E. R. et al. Implementation of the LandTrendr Algorithm on Google Earth Engine. Remote Sensing 10, doi:10.3390/rs10050691 (2018).

102 Gorsevski, V., Geores, M. & Kasischke, E. Human dimensions of land use and land cover change related to civil unrest in the Imatong Mountains of South Sudan. Appl. Geogr. 38, 64–75, doi:10.1016/j.apgeog.2012.11.019 (2013).

103 Kennedy, R. E., Yang, Z. & Cohen, W. B. Detecting trends in forest disturbance and recovery using yearly Landsat time series: 1. LandTrendr—Temporal segmentation algorithms. Remote Sens. Environ. 114, 2897–2910 (2010).

104 Griffiths, P., Van Der Linden, S., Kuemmerle, T. & Hostert, P. A pixel-based landsat compositing algorithm for large area land cover mapping IEEE Journal of Selected Topics in Applied Earth Observations and Remote Sensing 6, 2088–2101 (2013).

105 Cohen, W. B. & Spies, T. A. Estimating structural attributes of Douglas-fir/western hemlock forest stands from Landsat and SPOT imagery. Remote Sens. Environ. 41, 1–17 (1992).

106 Czerwinski, C. J., King, D. J. & Mitchell, S. W. Mapping forest growth and decline in a temperate mixed forest using temporal trend analysis of Landsat imagery, 1987–2010. Remote Sens. Environ. 141, 188–200 (2014).

107 Cohen, W. B., Yang, Z. & Kennedy, R. Detecting trends in forest disturbance and recovery using yearly Landsat time series: 2. TimeSync—Tools for calibration and validation. Remote Sens. Environ. 114, 2911–2924 (2010).

108 Grogan, K., Pflugmacher, D., Hostert, P., Kennedy, R. & Fensholt, R. Cross-border forest disturbance and the role of natural rubber in mainland Southeast Asia using annual Landsat time series. Remote Sens. Environ. 169, 438–453 (2015).

